# Active field theory approach to explain size control of transcriptional condensates

**DOI:** 10.64898/2026.05.17.725716

**Authors:** Kathrin Hertäg, Samuel Shoup, Leonhard T. Thews, Radhika Khatter, Eugenio Ferrario, Joshua F. Robinson, Sina Wittmann, Sandra Schick, Thomas Speck

## Abstract

Transcription factors organize into liquid-like condensates to facilitate gene expression, yet the physical mechanisms governing their formation and properties remain poorly understood. We study the size statistics of transcriptional condensates in human HAP1 cells using widefield and super-resolution microscopy tagging the epigenetic reader BRD4. We find that hubs that appear monolithic in widefield resolve into clusters of smaller droplets that resist coarsening. We link this size control to Active Model B+, a non-equilibrium field theory that captures a regime of reverse Ostwald ripening out of thermal equilibrium. In this regime, chemically driven currents cause larger droplets to transfer mass back to smaller ones, stabilizing a state of microphase segregation. The observed exponential size distribution of BRD4 foci quantitatively matches our numerical simulations, suggesting a universal physical picture for the non-equilibrium self-limitation of cellular condensates.

## 1. Introduction

Historically, the understanding of eukaryotic transcription by RNA polymerase II has been rooted in models depicting the sequential and hierarchical assembly of stable macromolecular complexes on DNA [1]. These models often emphasize high-affinity, lock-and-key type interactions between transcription factors (TF) and their cognate DNA sequences, followed by the recruitment of co-activators, chromatin remodelers, and the core transcriptional machinery. However, the idea of long-lived, stably bound transcriptional complexes is challenged by live-cell imaging and advanced genomic techniques [2, 3], which reveal a highly dynamic and transient nature of TF-chromatin interactions [4, 5]. Current models have thus shifted away from this static picture towards *transcription condensates* as dynamic, liquid-like environments that concentrate the necessary factors for efficient transcription at specific genomic locations such as enhancers and promoters [6, 7].

The ability of liquid-like condensates to form is often rationalized through the concept of phase segregation driven by reversible (weak) multivalent attractive interactions that overcome the entropic cost of concentrating molecules [8, 9] and are modulated by the environment (e.g., chromatin [10, 11] and salt concentration [12]). In the most simple physical picture–often compared to the immiscibility of oil in water–an initial distribution of droplets undergoes “coarsening”, through which the system evolves toward a single macroscopic droplet to minimize its interfacial area and free energy. This evolution is driven by conventional *Ostwald ripening* and larger droplets grow at the expense of smaller ones [13, 14].

At variance with this picture, cellular condensates appear to resist coarsening and maintain their size over long times. This resistance to coarsening presents a significant challenge to our physical understanding, particularly because these condensates occur in highly heterogeneous crowded environments and in the presence of chemically driven processes. Various microscopic mechanisms have been suggested to account for this behavior, including (but not limited to) the viscoelastic nature of the cellular environment [15–19] and “surfactant” effects from protein diversity that lower interfacial tension [20]. Quite generally, any competition between short-range and long-range interactions is sufficient to induce microphase separation and patterns [21, 22]. While thermodynamic models can explain some aspects of self-limiting assembly [23], the lack of geometric termination motifs and observations such as the rapid aging of condensates on ATP depletion [24] challenge conventional equilibrium theories and the role of chemically driven processes that impact protein interactions has moved into the focus [25, 26].

Still, most current theoretical approaches are rooted in equilibrium concepts and based on the physics of free-energy minimization. In contrast, the cell hosts a multitude of active processes that are driven through various means such as external chemical gradients, ATP hydrolysis, etc. Resolving these microscopic processes in an environment as complex as that of a living cell, and incorporating them into thermodynamically consistent models is an enormous challenge. Recent progress has shown how to include chemically driven modifications such as phosphorylation into coarse-grained models [27], which constitute an important computational tool for the study of biological condensates [28]. Instead of resolving single molecules and their detailed interactions, top-down frameworks constitute complementary approaches that model the evolution of continuous concentration fields and allow to shift the focus to the role of collective quantities such as the interfacial tension and their extensions to non-equilibrium [29]. Traditional frameworks include Flory-Huggins and Model B, which successfully describe passive phase segregation. Non-equilibrium then manifests itself through terms that cannot be derived from a free energy. A paradigmatic example is the extension of Model B to “Active Model B+” through higher-order gradient terms [30, 31]. Strikingly, for certain parameters the conventional Ostwald ripening scenario is replaced by the reverse process, whereby molecules diffuse from larger to smaller droplets.

In this manuscript, we explore to what extent the scenario of *reverse Ostwald ripening* is compatible with the behavior of condensates involving the bromodomain-containing protein 4 (BRD4). BRD4 is a key epigenetic regulator of transcription and is enriched at super-enhancers, dense clusters of regulatory elements that drive high levels of transcription, especially of cell identity and cell-type specific genes [32]. While BRD4 undergoes passive phase segregation in vitro, in cellulo super-resolution microscopy data reveals that droplets remain stable over hours without coarsening. By analyzing the size distribution of these foci and comparing them to numerical simulations, we link these biological transcription hubs to reverse Ostwald ripening as active non-equilibrium mechanism for size control.

## 2. Results and discussion

### 2.1. Transcription condensates in vitro and in cellulo

The epigenetic reader BRD4 contains two bromodomains that recognize and bind to acetylated lysine residues on histone tails [33]. Through this interaction, BRD4 is recruited to active enhancers within euchromatin as these are enriched in acetylated histone marks and thus binds independent of the DNA sequence. BRD4 functions as a scaffold that recruits P-TEFb (CDK9/cyclin T) to promoter regions, enabling CDK9-mediated phosphorylation of the RNA polymerase II C-terminal domain and negative elongation factors, thereby releasing RNA polymerase II from promoter-proximal pausing [34]. The ability of BRD4 to drive the formation of transcription hubs has been linked to its intrinsically disordered regions (IDRs) [35]. BRD4 exists primarily as two major isoforms, long (BRD4-L) and short (BRD4-S), which possess distinct C-terminal domains that lead to different functional roles [36].

Although phase segregation of BRD4 has been reported in vitro [35], quantitative data and phase diagrams are rare. To fill this gap, we performed a series of in vitro experiments using the long isoform (BRD4-L) varying temperature and (global) concentration. The results are shown in Figure 1A-C (and Supplemental Figure 1). BRD4 droplets were formed in reaction tubes and immediately encapsulated in oil-emulsion droplets. After centrifugation to facilitate the coalescence of droplets, these were imaged to estimate the total volume fraction [37]. Exploiting the lever rule, the linear extrapolation then yields the coexisting dilute (c−) and dense (c_+_) concentrations outside and inside the droplet, respectively (Figure 1A). For all considered temperatures between 10 °C and 40 °C we observe phase segregation, for which the coexistence region narrows when going to larger temperatures as expected for an upper critical solution temperature. For the dilute background concentration, we find c− = 2 µM, which appears to be independent of temperature (Figure 1B). While extrapolation to the dense concentrations c_+_ depends on a rather small range of global concentrations, the concentration c_+_ inside droplets is in the range of a few mM (Figure 1C) and thus three orders of magnitude larger than c−.

**FIGURE 1:**
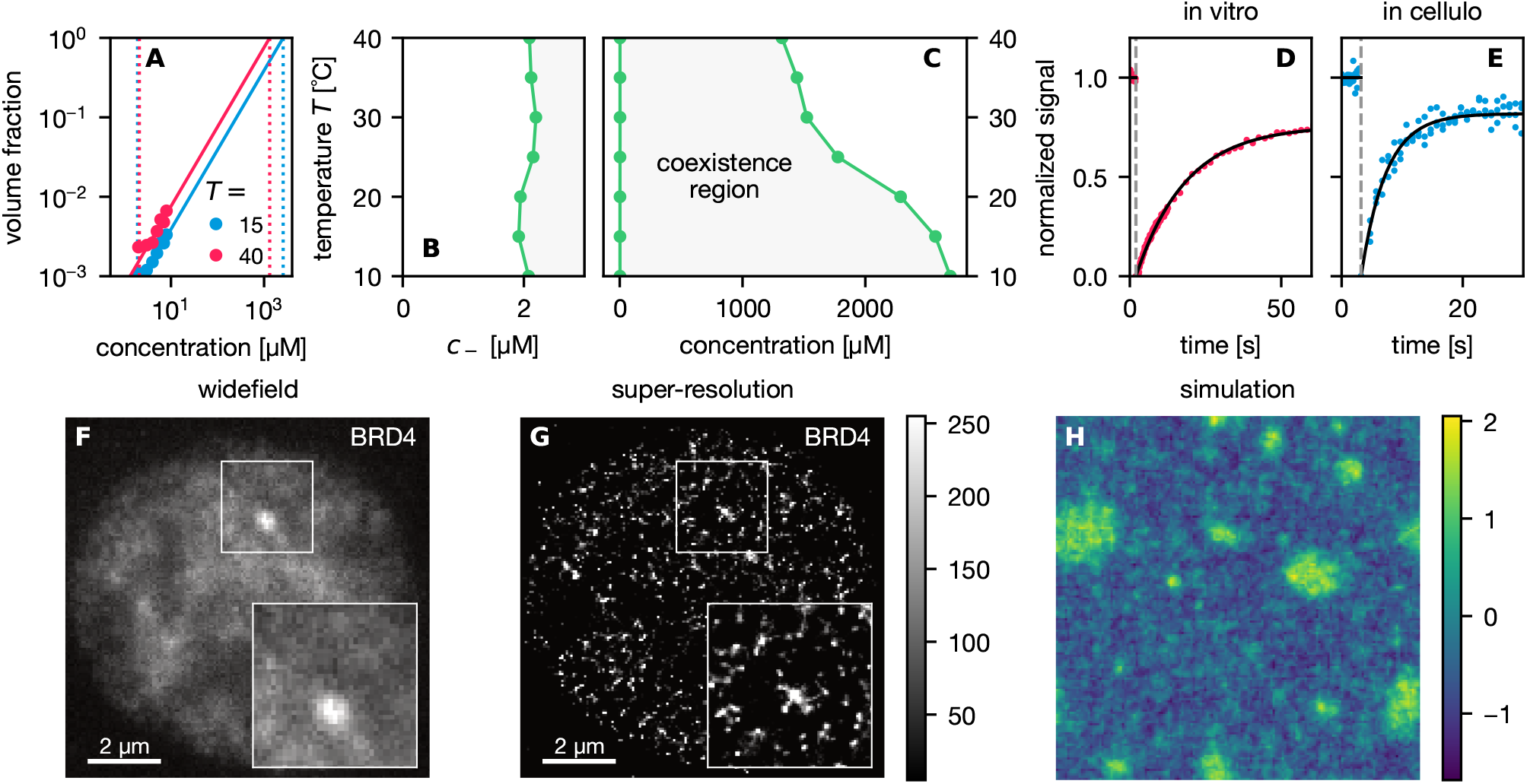
Phase segregation of BRD4. (A-C) In vitro results showing (A) measured volume fractions as function of total concentration for two temperatures T = 15 °C and T = 40 °C. Dashed lines are the extrapolated concentrations c*±*at which the volume fraction is zero and one, respectively. (B) Zoom onto the dilute binodal c− (T) of (C) the full phase diagram. (D,E) Signal recovery after FRAP bleaching (dashed line) can be fitted by a single exponential (solid line) for both (D) in vitro and (E) in cellulo. (F,G) Representative microscopy images obtained through (F) widefield and (G) super-resolution microscopy in human HAP1 cells of fluorescently tagged BRD4 imaging the same nucleus. While the widefield image exhibits a single focus, this focus resolves into smaller droplets at higher resolution. (H) Simulation snapshots of Active Model B+ (ζ =− 4 and α = −3.8) demonstrating fluctuating droplets of finite size in steady state. The colors indicate the value φ of the dimensionless field representing the relative concentration of the tagged molecules.

To probe the dynamic behavior of BRD4 in droplets, we perform fluorescence recovery after photobleaching (FRAP) experiments [38]. Bleaching a circular spot with radius R = 1 µm larger than the droplet (full FRAP), the recovery of the fluorescence signal follows a single exponential with relaxation time τ (Figure 1D and Supplemental Figure 2). Such a rapid recovery indicates high molecular mobility of BRD4 and a dynamic exchange of molecules between the droplet and the surrounding dilute environment through the interface of the droplet.

In the next step, BRD4 is tagged and monitored in living cells [39] using human HAP1 cells [40]. Typical widefield or confocal microscopy images show a few dispersed foci per nucleus (Figure 1F, Supplemental Figure 3). Remarkably, these foci appear to be stable over hours without showing coarsening and they resolve before mitosis and quickly reform after (Supplemental Figure 3). The size of these foci is close to the lateral resolution limit of confocal microscopy with an area of a few pixels, which corresponds to foci radii of approx. 250 nm. To provide a more detailed picture, we have gathered super-resolution microscopy images, which now exhibit a wide range of foci sizes below the resolution limit of confocal microscopy (Figure 1G). In particular, the foci seen in widefield microscopy resolve into smaller droplets.

To investigate the dynamic response of foci, we again performed FRAP experiments with bleaching areas larger than foci. As shown in Figure 1E, the recovery follows a single exponential as in the in vitro experiments (Figure 1D). While we refrain from a quantitative comparison (in particular due to the presence of both isoforms in cells and the fact that BRD4 is expected to be predominantly bound to chromatin) we conclude that foci remain dynamic, and that there is an exchange of tagged BRD4 molecules between foci and the nucleoplasm. Importantly, the observed suppression of coarsening is not due to a kinetic arrest, calling for a different physical mechanism underlying size control in transcription condensates.

### 2.2. Theoretical insights into size control

Before discussing the statistics of foci sizes and their implications, we briefly review theoretical approaches of phase segregation. The basic phenomenon of phase segregation into domains exhibiting different compositions is already captured by simple models that encapsulate the competition between the gain of (free) energy due to attractive interactions and the loss of translational entropy, the application of which to biomolecular condensates has been reviewed extensively (e.g., Refs. 8, 9). In a nutshell, for a single tagged component with local concentration c, the free energy density f(c) has a generic shape that develops two local minima. The coexisting concentrations c*±*inside the different domains then correspond to equal slope of the free energy density (Figure 2A), i.e., equal chemical potential µ. The interface between regions of low and high concentration is mainly governed by the interfacial tension. In this simple picture, an initial distribution of droplets “coarsens” and the system evolves towards a single macroscopic droplet (as seen in the in vitro experiments of BRD4). This evolution is driven by the minimization of the interfacial area and the free energy cost associated with it. One molecular mechanism is (conventional) Ostwald ripening driven by droplet curvature, moving molecules from small droplets with high curvature to big droplets with small curvature along the gradient of the chemical potential µ until the small droplets have vanished and µ has attained a uniform value. In other words, the chemical potential µ(R) close to droplets is a function of their radius R so that µ(R) is a decreasing function and smaller radii have a larger value (Figure 2B).

**FIGURE 2:**
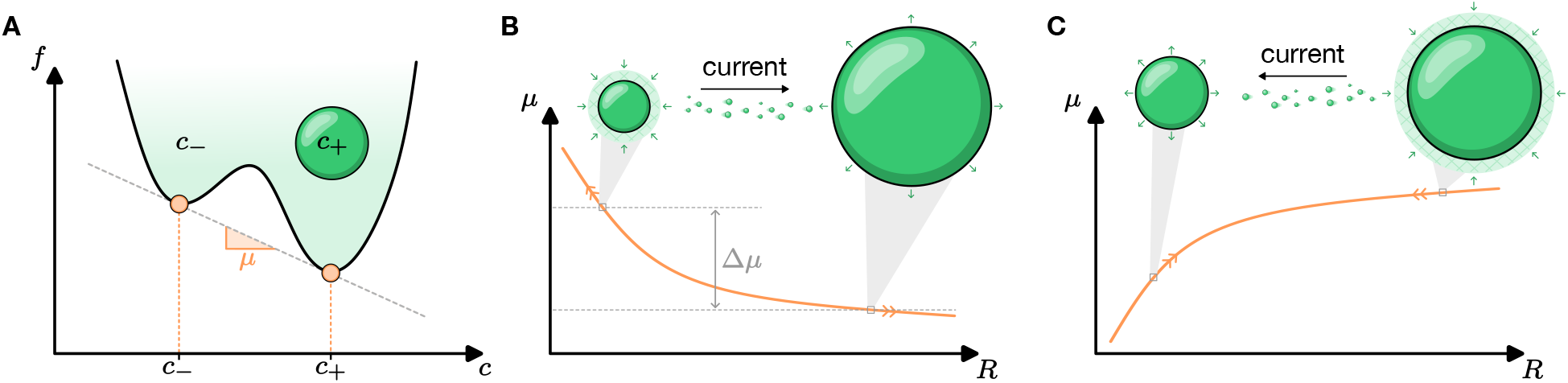
Theory of phase segregation. (A) Sketch of a typical free energy f(c) depending on the concentration c, which exhibits two minima. In thermal equilibrium, the coexisting concentrations c*±* are determined from the common tangent with slope µ corresponding to a uniform chemical potential. (B,C) Ripening is driven by the gradient of the chemical potential. (B) In the case of conventional Ostwald ripening, smaller droplets have a larger chemical potential close to their surface and the current transports molecules from small to large droplets, leading to a single macroscopically big droplet at coexistence with a dilute environment. (C) Away from thermal equilibrium we assume that the current is still be governed by an effective chemical potential. It’s dependence on the droplet radius, however, might be reversed, leading to a current that now stabilizes smaller droplets at the expense of larger ones.

Clearly, the situation in cellulo is far more complicated with segregation occurring in a dense crowded environment and in the presence of chemically driven processes. The foci shown in Figure 1F and G resist coarsening over hours (Supplemental Figure 3). As briefly reviewed in the introduction, this behavior has been rationalized through a range of underlying microscopic mechanisms without reaching a consensus yet. All these approaches focus on a specific physical mechanisms. A more agnostic and physics-inspired approach is to ask for “universality”, an overarching class into which these models fall and that captures the basic phenomena without making detailed assumptions on the underlying molecular processes. For instance, the passive phase segregation captured by a single field described above is known as “Model B” according to the famous classification proposed by Hohenberg and Halperin [41]. It captures a wide range of phenomena in systems as diverse as binary metallic alloys, polymer blends, and the spinodal decomposition in silicate glasses. Crucially, such theories describe the limiting behavior at large scales, at which most details of the interactions get subsumed into a few effective coefficients of the theory.

While Model B encapsulates the physics of short-range interactions, additional long-range interactions allow for microphase segregation with a host of regular patterns (lamellar, hexagonal, etc.) in thermal equilibrium [21], including regular arrangements of finite droplets. Crucially, these patterns are governed by an explicit length scale that can be identified through linear stability analysis.

Recently, Model B has been extended by additional terms that cannot be derived from a free energy, but that appear in an expansion of gradients [30, 42], see Section 4 for further details. These new terms come with additional coefficients that quantify their influence, which we denote α and ζ in the following. These coefficients encode microscopic interactions but cannot be assigned directly to single specific molecular properties (comparable in spirit to the χ-parameters of Flory-Huggins theory). Remarkably, this “Active Model B+” captures a host of phenomena that divert from the passive physics of Model B and that have come to be associated with biomolecular condensates. In particular, there are now regions in parameter space where coarsening is replaced by a novel mechanism that has been coined “reverse Ostwald ripening” and that suggests a physical large-scale mechanism by which size control of droplets can be achieved [30, 43]. Figure 1H shows a snapshot from numerical simulations of Active Model B+. It demonstrates that coarsening is indeed suppressed and the steady state exhibits a range of finite-size droplets that qualitatively resemble the super-resolution images.

Importantly, Active Model B+ does not simply introduce a length scale that sets the preferred droplet distance since the linearized current [Eq. (2)] is exactly the same as passive Model B. In Figure 2C we sketch how reverse Ostwald ripening works in Active Model B+ without an external length scale. Even without a free energy, we can still define a chemical potential that governs currents [42, 44]. In contrast to Model B, however, for some values of the coefficients (ζ, α) the function µ(R) switches sign and increases for larger droplets, implying that the current reverses and molecules move from larger to smaller droplets. Considering two droplets of different sizes, the large droplet thus transfers molecules to its environment and the smaller droplet, which grows until both have reached the same size and the chemical potential is uniform throughout the system. In the presence of noise we then observe a highly dynamic steady state typically associated with “microphase” segregation, in which droplets nucleate, grow, and shrink, they sometimes even merge or split, but they do not coalesce into a single domain [42, 45]. Importantly, the growth of droplets is not kinetically arrested and molecules diffusively move into and out of droplets.

### 2.3. Statistics of foci sizes

In Figure 3A, we plot the histogram of foci areas obtained from the super-resolution microscopy images processing 389 foci from 22 nuclei. The area is measured in pixels with a lower cutoff of 10 pixels. The histogram shows a nearly exponential decay for smaller sizes with three outliers at larger areas (with radii of approx. 160 nm) separated by a gap. Converting the area of 150 pixels implies a radius of 160 nm for the large foci, which at a concentration of approx. 2 mM (cf. Figure 1C) would contain on the order of 10^4^ molecules. Such a distribution of cluster sizes is qualitatively compatible with the formation of finite micelles. However, assuming that BRD4 is indeed driving the formation of condensates, and taking into account that we have established that in vitro BRD4 undergoes phase separation, micellization of such a large number of molecules seems unlikely.

**FIGURE 3:**
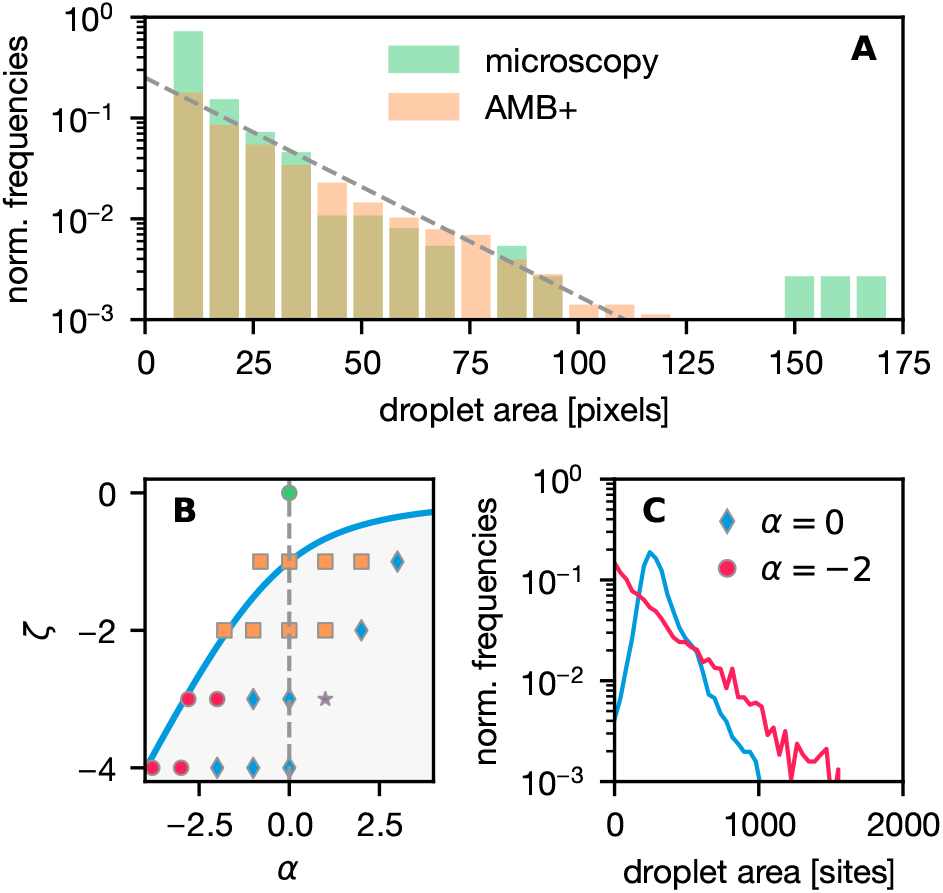
Droplet statistics and mapping to model. (A) Histogram of droplet areas obtained from super-resolution microscopy overlaid by the simulation results (ζ = −4 and α = −3). (B) Phase diagram of Active Model B+ spanned by the two coefficients ζ and α. The solid line is the mean-field prediction for the transition from conventional to reverse Ostwald ripening (within shaded area). The symbols indicate: conventional macrophase segregation (□), microphase segregation (⋄and◦), and a phase of spatially ordered droplets (⋆). The passive in vitro experiments sit at the origin (green symbol). (C) Two histograms at ζ = −3 of simulated droplets demonstrating the difference within the region of microphase separation.

Assuming that the outliers are rather due to finite statistics, in Figure 3A we also show the distribution of (rescaled) areas obtain from numerically solving Active Model B+ on a two-dimensional grid. Instead of the concentrations c, in the model we consider a rescaled dimensionless field *ϕ* so that in the passive limit *ϕ*_−_= −1 indicates the dilute environment and *ϕ*_+_ = +1 the inside of dense droplets. The mapping to concentrations thus is 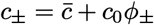with concentrations 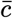(also known as rectilinear diameter) and amplitude c_0_. In the reverse Ostwald ripening regime, the concentration inside droplets increases while the background concentration decreases as the droplets become smaller. This effect can be seen in Figure 1H (corresponding to ζ = − 4 and α = − 3.8), where field values inside droplets reach *ϕ*_+_ ≃ 2 compared to the passive values *ϕ*_±_ ≃ ± 1 for the corresponding Model B (same parameters except ζ = α = 0).

Figure 3B shows the phase diagram of Active Model B+ in the plane spanned by ζ and α holding all other model parameters fixed [42]. We show the mean-field prediction for the transition from conventional to reverse Ostwald ripening together with the results of “noisy” simulations (see Section 4 and Ref. 42 for details). The simulations find conventional coarsening into a single droplet even within the regime where microphase segregation is predicted in the absence of noise. The region wherein simulations confirm the steady-state existence of multiple droplets can be further divided into three subclasses: a regular dense lattice of droplets (which is of no interest here), distributions of sizes with a peak and fast decay, and, moving close to the transition line, distributions characterized by the vanishing of the peak accompanied by a slower exponential decay (Figure 3C). The later distributions agree well with the experimental super-resolution data as shown in Figure 3A.

## 3. Conclusions

Uncovering the physical mechanisms behind size control of protein condensates in cellular environments is an open challenge. Here we have extracted the size statistics of BRD4 foci associated with transcription condensates in living cells using super-resolution microscopy. While widefield microscopy has supported the picture of a few isolated stable condensates, these resolve into clusters of neighboring droplets (the largest of which reach radii of approx. 160 nm) that optically fuse in widefield. Outside these clusters many more foci are observed with a wide distribution of sizes that follows an exponential decay. Such an exponential decay implies the lack of both a typical size of and typical distance between droplets. In thermal equilibrium, it would require a free energy that scales exactly with the number of molecules (here up to ∼10^4^). While possible for linear aggregates [46], the different scaling of droplet volume and area rules out such an explanation. Microemulsions, micelles, and other self-limited equilibrium structures exhibit size distributions that are peaked around a typical size. Our observations (including the dynamic exchange of proteins between droplet and nucleoplasm) thus point to dynamic liquid-like behavior of BRD4 that is at least compatible with Active Model B+, a field theory for active liquids. Matching with detailed simulation results, we propose that condensates reside close to the transition from macrophase to microphase separation for large negative values of the coefficient ζ. The implication is that condensates are actively maintained away from thermal equilibrium, and the physical mechanism behind size control is reverse Ostwald ripening. Our results constrain microscopic models and give a first glimpse into potential universality underlying cellular condensates.

## 4. Experimental section

### 4.1. Protein expression and purification

A recombinant Twin-Strep–MBP–3Csite-mEGFP-HsBRD4-L fusion was expressed in Sf9 insect cells using the Bac-to-Bac system (Invitrogen). Cells were infected at a density of approximately 1 ×10^6^ cells/mL and cultured in SF900 III medium (Gibco) at 27 °C for 55 h prior to harvest. Cell pellets were collected by centrifugation (250 ×g, 15 min, 8 °C), resuspended in ice-cold lysis buffer (50 mM Tris-HCl, 500 mM NaCl, 5% glycerol, 0.2% Triton X-100, 2 mM EDTA, 2 mM mM DTT, Protease Inhibitor Cocktail III (EMD Millipore, pH 7.5), and lysed by sonication. Lysates were clarified by centrifugation (55, 000× g, 30 min, 4 °C) and incubated with Strep-Tactin XT resin (IBA Lifesciences GmbH) for 1–16 h for affinity capture. The resin was washed extensively with wash buffer (50 mM Tris-HCl, 500 mM NaCl, 5% glycerol, pH 7.5), including a high-salt wash step (wash buffer with 2 M NaCl). Bound protein was released by on-resin cleavage with 3C protease (33 µg mL^−1^, produced in-house) in the presence of 2 mM DTT for 10 min at room temperature. Cleaved BRD4-L was further purified by size-exclusion chromatography on a Superdex 200 Increase 16/600 column equilibrated in gel filtration buffer (25 mM HEPES, 400 mM NaCl, 5% glycerol, 1 mM DTT, pH 7.4). Peak fractions were pooled and concentrated using a 30 kDa molecular weight cutoff Amicon concentrator (Merck Millipore), snap-frozen in aliquots, and stored at -80 °C until use.

### 4.2. In vitro phase diagram

Phase diagrams were generated using the “inPhase” method as described by Fritsch *et al*. [37]. In brief, purified BRD4-L was thawed on ice and cleared by centrifugation (10 min, 21, 000 ×g, 4 °C) prior to use. BRD4-L was diluted at 10 °C to the desired final concentrations, yielding samples with a final buffer composition of 15 mM HEPES (pH 7.4), 50 mM NaCl, 0.5% glycerol, 1 mM DTT, and 1 mM EDTA. Samples were immediately mixed 1:1 (v/v) with pre-cooled fluorinated Pico-Surf oil (Sphere Fluidics) to generate emulsion droplets. Emulsions were centrifuged briefly (3 min, 200× g, 10 °C) and loaded into custom-made channels integrated into a temperature-controlled Vulcan slide stage (Blue Ice Labs GmbH) set to 10 °C. Temperature was increased in 5 °C increments with 10 min equilibration at each step prior to imaging.

Imaging was performed on a Leica THUNDER wide-field microscope (Leica Microsystems) controlled with LAS X software. Samples were imaged using a 20 ×objective (NA 0.80). Brightfield images and fluorescence z-stacks (1 µm step size) spanning all condensate droplet heights were acquired. mEGFP was excited using an LED light source at 488 nm (LED8 by Leica) and a DFT51010 quadband filter cube (Leica) and 535/70 emission bandpass filter were used to split fluorescence emission from excitation light. Images were recorded over an exposure time of 10 ms (brightfield) and 5 ms (mEGFP fluorescence) on an K8 sCMOS camera (Leica) that was operated with no binning, generating images with a pixel size of 325 nm in the object plane. Fluorescence maximum-intensity projections were analyzed in Fiji/ImageJ using a custom script [47, 48]. Individual emulsion droplets were defined based on the brightfield image by manual regions of interest (ROIs), and droplet volumes were estimated from ROI dimensions assuming spherical geometry, with a maximum height set by the channel height (100 µm). Condensates were segmented within each emulsion droplet in the fluorescence maximum projection, and the dense-phase volume was estimated by summing spherical volumes derived from the segmented condensate areas. The volume fraction was calculated as the ratio of dense-phase volume to emulsion droplet volume for each emulsion droplet.

### 4.3. In vitro FRAP

Imaging of GFP-BRD4-L droplets in vitro was performed using Leica TCS SP5 STED confocal microscope (Leica Microsystems) with a 40× oil immersion objective. Droplets were formed by mixing 4 µM BRD4-L in a buffer containing 15 mM Na-Hepes, 50 mM NaCl, 0.5% glycerol, 1 mM DTT, 1 mM EDTA. After incubation for 5 min, imaging was performed on droplets that had settled close to the imaging surface.

For photobleaching, a 1.5 µm diameter ROI was used for spot bleach, and entire droplets with approx. 2 µm diameter were used for full bleach. Bleaching was carried out with 488 nm laser at 70% power for 5 frames at

0.099 s per frame. Pre-bleach images were acquired for 20 frames at 0.099 s/frame. Post-bleach images for spot bleach were collected for 50 frames at 0.09 s/frame, then 50 frames at 2 s/frame and for full bleach for 100 frames at 0.09 s/frame, followed by 50 frames at 2 s/frame. Fluorescence recovery for spot bleach was quantified using EasyFRAP with double normalization [49].

### 4.4. Cell culture

Human HAP1 cells [50, 51] were genetically engineered with CRISPR/Cas9 to express intron-tagged BRD4 with eGFP (kindly provided by Stefan Kubicek) and MED14 or POLR2A C-terminally-tagged with TagRFP [40].

HAP1 cells were maintained at 37 °C in a humidified incubator (5% CO_2_, 85% relative humidity) and cultured in Iscove’s Modified Dulbecco’s Medium (IMDM) with phenol red supplemented with 10% (v/v) fetal bovine serum (FBS) without antibiotics. Cells were passaged approximately twice per week at 70-80% confluence. For passaging, medium was aspirated, the plate rinsed once with DPBS, and cells detached with 0.25% trypsin/EDTA for 5 min at 37 °C. Trypsinization was neutralized with 4 mL medium; cells were collected, pelleted (300 ×g, 5 min), resuspended in fresh warm medium, and reseeded at the required density (typically 1:12 diluted for regular passaging) into 10 cm dishes filled with 10 mL fresh medium.

For imaging experiments, cells were seeded 24 h prior to imaging. Poly-D-lysine (PDL) coating was used to increase cell spreading for 96-well and 384-well plates: 50 µL PDL (50 µg mL^−1^ in DPBS) per well was incubated for one hour at room temperature (RT) on an orbital shaker (30 rpm). PDL solution was removed and 200 µL DPBS added for plate storage at 4 °C (≤2 weeks). Before use, DPBS was removed and plates air-dried for one to three hours with the lid open in a laminar flow cabinet. Harvested cell suspension densities were measured following spin-down and resuspension in fresh medium using an automated cell counter in brightfield mode. Cells were seeded at 20,000 cells per well in 200 µL medium for 96-well plates or 5,000 cells per well in 50 µL medium for 384-well plates.

### 4.5. Live-cell confocal microscopy

Imaging was performed on an Opera Phenix High-Content Screening System (Revvity) in spinning-disk confocal mode (Harmony software v.5.2). The environmental chamber was maintained at 37 °C and 5% CO_2_ (pre-heated). Prior to imaging, Hoechst 33342 nuclear staining was performed: medium was removed and replaced with 50 µL labeling solution (1 µg mL^−1^ Hoechst 33342 in IMDM). Labeling lasted for 30 minutes in the incubator under standard conditions and was followed by three 200 µL brief wash steps in warm medium. The final wash step was left on cells for 30 minutes under standard conditions to allow unbound dye to diffuse out of cells.

HAP1 cells were imaged in 96-well multiwell plates (PhenoPlate, Revvity) in warm phenol-red-free IMDM, which was exchanged before imaging. Images were acquired with flatfield correction and maximum intensity projections created from z-stacks. Laser excitation, emission, exposure, and power were kept constant: Hoechst 33342 (405 nm excitation, 435-480 nm emission, 120 ms exposure, 60% power); eGFP/BRD4 (488 nm excitation, 500-550 nm emission, 700 ms exposure, 70% power); TagRFP/MED14 or POLR2A (561 nm excitation, 570- 630 nm emission, 700 ms exposure, 70% power). For each field of view, z-stacks covering the full nuclear volume (step size 0.3-0.5 µm) were acquired.

### 4.6. Live-cell FRAP

FRAP was used to quantify BRD4::eGFP exchange within nuclear spots by measuring fluorescence recovery after targeted bleaching. HAP1 BRD4::eGFP MED14::TagRFP cells were harvested normally, counted, and seeded at 75,000 cells per well in 8- well µ-Slide chamber slides in 300 µL medium. After 24 hours, cells were maintained at 37 °C, 5% CO_2_ during imaging.

Imaging was performed on a spinning-disk confocal microscope (W1-disk, 50 µm pinhole, Yokogawa) based on a Ti2-E microscope (Nikon) with a FRAP unit (Visitron Systems) controlled by VisiView software. FRAP was performed using a 60× water immersion objective (NA 1.2, CFI plan apo VC), 2×2 binning, and 488 nm excitation for GFP (300 ms exposure). Photobleaching was executed with the 488 nm FRAP path. For each cell, a fixed circular region of interest (ROI) of three-pixel radius centered on a BRD4–GFP focus was bleached. Acquisition sequence: (1) Pre-bleach: 10 frames at maximum frame rate (300 ms each); (2) Bleach: 5 bursts (no imaging during bleach; ROI radius 3 pixels; 100 ms/pixel dwell time); (3) Recovery: 60 frames at 1 frame/s.

Series were processed in Fiji (ImageJ 2.9.0/1.53t) with a custom macro to extract mean intensities for the bleached ROI, a manually selected background region, and nuclear region. Traces were normalized in easyFRAP using the full-scale method (background/reference correction) and fit with a single-exponential model to estimate mobile fraction and half-time of recovery (t_1*/*2_). Traces were excluded if the ROI/nucleus moved out of frame or if fit quality was poor.

### 4.7. Super-resolution microscopy

HAP1 BRD4::eGFP POLR2A::TagRFP cells were seeded in 8-well chamber slides (75, 000 cells/well, 300 µl medium) and cultured for 24 hours under standard conditions. Cells were fixed with 4% paraformaldehyde (PFA) in DPBS (pre-warmed to 37°C) for 30 minutes at RT. Following 5-minute incubation in immunofluorescence (IF) quenching buffer (50 mM Tris, 100 mM NaCl in PBS) on a slow-moving orbital shaker, cells were treated with permeabilization buffer (0.2% Triton X-100 in PBS) for 20 minutes, washed 2×with PBS, and blocked in 3% bovine serum albumin (BSA) in PBS for 30 minutes at RT. 100 µl of antibody solution (1 : 500 FluoTag-Q anti-GFP conjugated to Alexa Fluor 647 in 1% BSA in PBS) was added and incubated for 1 hour (RT, dark). Following 4 ×wash in PBS, samples were stored at 2-8 °C in PBS until imaging.

Immediately prior to imaging, PBS was replaced with GLOX/β-mercaptoethanol (BME) switching buffer prepared in 50 mM Tris-HCl, 10 mM NaCl (pH∼8) and containing glucose oxidase/catalase (GLOX/CAT), 10% (w/v) glucose, and 50 mM β-mercaptoethanol. Working 5 mL mixes were assembled from stocks as follows: 50 µl GLOX (100 ×stock), 50 µl catalase (100 ×stock), 500 µl glucose (10× stock), 50.3 µl of 1 M β-mercaptoethanol (final 50 mM), and Tris-HCl buffer to volume.

Single-molecule localization microscopy (SMLM) was performed on a NanoImager microscope (ONI) in widefield mode using 647 nm excitation for Alexa Fluor 647. Imaging was controlled with NimOS vendor software. Emission was collected through standard AF647 filter sets. For each field, a brief widefield reference image (no blinking) was recorded, then SMLM movies were acquired for 20,000 frames under continuous illumination. Raw microscopy images were processed using ImageJ/Fiji (ImageJ 2.9.0/1.53t) and the ThunderStorm plugin. Following drift correction and exclusion of the first 1,000 images, average shifted histogram visualizations were prepared using 5X pixel magnification. Localizations were obtained using a B-spline wavelet filter (order 3, scale 2.0), local-maximum detection (8- neighborhood, threshold 1.5×std(Wave.F1)), integrated-Gaussian point spread function with maximum likelihood estimation fitting (initial σ = 1.6 px, radius 3 px), camera: 117 nm/px, 1.71 electron readout noise, 0.46 electron/analog-to-digital unit, quantum efficiency 0.82, without electron multiplying gain. Frames > 1000 were kept, drift corrected by cross-correlation (10 bins), reactivations merged (30 nm, off-frames ≤ 1, 3 ≤ frames/molecule), quality-filtered (80–300 nm σ; xy-uncertainty < 30 nm), and rendered at 5× magnification.

### 4.8. Statistical analysis

We employ the same analysis pipeline to process pixel-based super-resolution images and simulation snapshots (cf. Figure 1G,H): First, we apply Otsu’s thresholding method to distinguish droplet and background pixels. Pixel clusters of less than five pixels are excluded due to background noise. We reconstruct the original brightness of the pixels in droplets and a gray scale image is constructed. This step is necessary to apply thresholding once more to obtain a binary classification of droplet and background pixels, whereby single background pixels inside a droplet are interpolated as droplet pixels. We then assign mutually connected droplet pixels to droplets. Each droplet pixel with one background pixel within its eight neighbors is treated as an edge pixel. Further, due to periodic boundary conditions in the simulation, the algorithm searches for droplets at the picture edge and if there is part of a droplet on opposite sides those droplets are unified as one. Finally, the size of the droplets is extracted in pixels.

### 4.9. Modeling condensates

We focus on a single species (here BRD4) and seek a model that captures the evolution of a corresponding scaled field *ϕ*(**r**, t) in two dimensions. We note that even in the presence of many interconvertible components, a similar scalar field theory emerges on large length and time scales when constrained by a conservation law [52]. Assuming conservation of mass implies the continuity equation

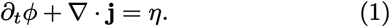

Following the general idea of Ginzburg-Landau and expanding the current **j** into gradients of the field implies a deterministic current [30]

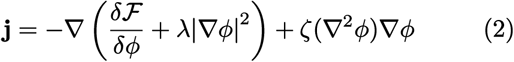

with (undetermined) coefficients λ and ζ through which the specific microscopic details of the system under scrutiny enter the model. In addition, Eq. (1) contains a Gaussian white noise field η(**r**, t) conserving *ϕ* with mean ⟨η(**r**, t)⟩ = 0 and correlations

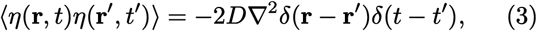

with the correlation strength set by D. The global “mass” is conserved with

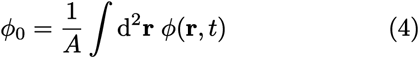

with area A in the system. The particle current involves the derivative of the Ginzburg-Landau free energy functional

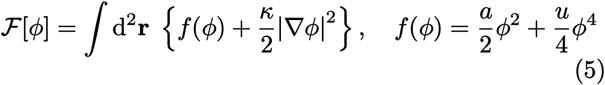

parametrized through the coefficients a, u; and κ penalizes inhomogeneities of the field. Inhomogeneous systems require *a* < 0 and passive Model B is recovered for ζ = λ= 0 with coexisting field values 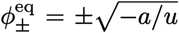 calculated as minima of the bulk free energy density *f*(*ϕ*).

We solve Eq. (1) numerically through spatially discretizing the field *ϕ* on a regular grid using periodic boundary conditions. Details can be found in Ref. 42. We measure lengths in units of the lattice spacing 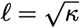 so that κ becomes unity in these units. Throughout, we fix u = −a = 0.25 and the noise strength D = 0.3 and report simulations for various values of ζ and α = ζ − 2λ.

## Acknowledgments

We acknowledge funding from the Deutsche Forschungsgemeinschaft (DFG) through CRC1551 “Polymer concepts in cellular function” (project number 464588647) and the CRC1551 support project Z01/IMB Protein Production CF for the assistance in generation of recombinant BRD4-L. We acknowledge support by the state of Baden-Württemberg through bwHPC and the German Research Foundation (DFG) through grant INST 35/1597-1 FUGG. SS was supported by the Max Planck Graduate Center with the Johannes Gutenberg-Universität Mainz (MPGC).

We thank Stefan Kubicek (CeMM, Vienna, Austria) for providing the HAP1 BRD4::eGFP cell line. We thank the IMB Microscopy and Histology CF team for their support in image acquisition and analysis, live-cell time-lapse imaging on the Visitron Spinning Disc (DFG project 402386039), high-throughput live-cell confocal imaging on the Opera Phenix (DFG project 316215830), FRAP on the TCS STED CW super-resolution microscope (DFG project number 204504610), and phase diagram aquisition with the THUNDER wide-filed microscope; and especially Márton Gelléri, Petri Turunen, and Anusha Gopalan for their assistance. We thank Juan M. Iglesias-Artola for advice adapting the inPhase protocol to BRD4.

## Data Availability Statement

The data that support the findings of this study are available from the corresponding author upon reasonable request.

## References

[1] G. Orphanides, T. Lagrange, and D. Reinberg, The general transcription factors of RNA polymerase II., Genes Dev. 10, 2657 (1996).

[2] I. I. Cisse, I. Izeddin, S. Z. Causse, L. Boudarene, A. Senecal, L. Muresan, C. Dugast-Darzacq, B. Hajj, M. Dahan, and X. Darzacq, Real-Time Dynamics of RNA Polymerase II Clustering in Live Human Cells, Science 341, 664 (2013).

[3] S. Chong, C. Dugast-Darzacq, Z. Liu, P. Dong, G. M. Dailey, C. Cattoglio, A. Heckert, S. Banala, L. Lavis, X. Darzacq, and R. Tjian, Imaging dynamic and selective low-complexity domain interactions that control gene transcription, Science 361, eaar2555 (2018).

[4] T. Mittag, L. E. Kay, and J. D. Forman-Kay, Protein dynamics and conformational disorder in molecular recognition, J. Mol. Recognit. 23, 105 (2010).

[5] K. Wagh, D. A. Stavreva, A. Upadhyaya, and G. L. Hager, Transcription Factor Dynamics: One Molecule at a Time, Annu. Rev. Cell Dev. Biol. 39, 277 (2023).

[6] D. Hnisz, K. Shrinivas, R. A. Young, A. K. Chakraborty, and P. A. Sharp, A Phase Separation Model for Transcriptional Control, Cell 169, 13 (2017).

[7] M. Palacio and D. J. Taatjes, Merging Established Mechanisms with New Insights: Condensates, Hubs, and the Regulation of RNA Polymerase II Transcription, J. Mol. Biol. 434, 167216 (2022).

[8] A. A. Hyman, C. A. Weber, and F. Jülicher, Liquid-Liquid Phase Separation in Biology, Annu. Rev. Cell Dev. Biol. 30, 39 (2014).

[9] S. F. Banani, H. O. Lee, A. A. Hyman, and M. K. Rosen, Biomolecular condensates: Organizers of cellular biochemistry, Nat. Rev. Mol. Cell Biol. 18, 285 (2017).

[10] J. A. Morin, S. Wittmann, S. Choubey, A. Klosin, S. Golfier, A. A. Hyman, F. Jülicher, and S. W. Grill, Sequence-dependent surface condensation of a pioneer transcription factor on DNA, Nat. Phys. 18, 271 (2022).

[11] A. R. Strom, J. M. Eeftens, Y. Polyachenko, C. J. Weaver, H.-F. Watanabe, D. Bracha, N. D. Orlovsky, C. C. Jumper, W. M. Jacobs, and C. P. Brangwynne, Interplay of condensation and chromatin binding underlies BRD4 targeting, MBoC 35, ar88 (2024).

[12] G. Krainer, T. J. Welsh, J. A. Joseph, J. R. Espinosa, S. Wittmann, E. de Csilléry, A. Sridhar, Z. Toprakcioglu, G. Gudiškytė, M. A. Czekalska, W. E. Arter, J. GuillénBoixet, T. M. Franzmann, S. Qamar, P. S. George-Hyslop, A. A. Hyman, R. Collepardo-Guevara, S. Alberti, and T. P. J. Knowles, Reentrant liquid condensate phase of proteins is stabilized by hydrophobic and non-ionic interactions, Nat. Commun. 12, 1085 (2021).

[13] P. W. Voorhees, The theory of Ostwald ripening, J. Stat. Phys. 38, 231 (1985).

[14] M. Cates, Complex fluids: The physics of emulsions, in Soft Interfaces, edited by L. Bocquet, D. Quéré, T. A. Witten, and L. F. Cugliandolo (Oxford University Press Oxford, 2017) 1st ed., pp. 317–358.

[15] R. W. Style, T. Sai, N. Fanelli, M. Ijavi, K. Smith-Mannschott, Q. Xu, L. A. Wilen, and E. R. Dufresne, Liquid-liquid phase separation in an elastic network, Phys. Rev. X 8, 011028 (2018).

[16] K. A. Rosowski, T. Sai, E. Vidal-Henriquez, D. Zwicker, R. W. Style, and E. R. Dufresne, Elastic ripening and inhibition of liquid–liquid phase separation, Nat. Phys. 16, 422 (2020).

[17] E. Vidal-Henriquez and D. Zwicker, Cavitation controls droplet sizes in elastic media, Proc. Natl. Acad. Sci. U.S.A. 118, e2102014118 (2021).

[18] H. Tanaka, Viscoelastic phase separation in biological cells, Commun. Phys. 5, 167 (2022).

[19] J. X. Liu, M. P. Haataja, A. Koŝmrlj, S. S. Datta, C. B. Arnold, and R. D. Priestley, Liquid–liquid phase separation within fibrillar networks, Nat. Commun. 14, 6085 (2023).

[20] A. W. Folkmann, A. Putnam, C. F. Lee, and G. Seydoux, Regulation of biomolecular condensates by interfacial protein clusters, Science 373, 1218 (2021).

[21] M. Seul and D. Andelman, Domain Shapes and Patterns: The Phenomenology of Modulated Phases, Science 267, 476 (1995).

[22] C. B. Muratov, Theory of domain patterns in systems with long-range interactions of Coulomb type, Phys. Rev. E 66, 066108 (2002).

[23] M. F. Hagan and G. M. Grason, Equilibrium mechanisms of self-limiting assembly, Rev. Mod. Phys. 93, 025008 (2021).

[24] A. Patel, H. O. Lee, L. Jawerth, S. Maharana, M. Jahnel, M. Y. Hein, S. Stoynov, J. Mahamid, S. Saha, T. M. Franzmann, A. Pozniakovski, I. Poser, N. Maghelli, L. A. Royer, M. Weigert, E. W. Myers, S. Grill, D. Drechsel,A. A. Hyman, and S. Alberti, A Liquid-to-Solid Phase Transition of the ALS Protein FUS Accelerated by Disease Mutation, Cell 162, 1066 (2015).

[25] D. Zwicker, The intertwined physics of active chemical reactions and phase separation, Current Opinion in Colloid & Interface Science 61, 101606 (2022).

[26] F. Jülicher and C. A. Weber, Droplet Physics and Intra-cellular Phase Separation, Annu. Rev. Condens. Matter Phys. 15, 237 (2024).

[27] E. Zippo, D. Dormann, T. Speck, and L. S. Stelzl, Molecular simulations of enzymatic phosphorylation of disordered proteins and their condensates, Nat. Commun. 16, 4649 (2025).

[28] J.-E. Shea, R. B. Best, and J. Mittal, Physics-based computational and theoretical approaches to intrinsically disordered proteins, Curr. Opin. Struct. Biol. 67, 219 (2021).

[29] H. Weyer, T. A. Roth, and E. Frey, Protein pattern morphology and dynamics emerging from effective interfacial tension, Nat. Phys. 10.1038/s41567-025-03101-6 (2025).

[30] E. Tjhung, C. Nardini, and M. E. Cates, Cluster Phases and Bubbly Phase Separation in Active Fluids: Reversal of the Ostwald Process, Phys. Rev. X 8, 031080 (2018).

[31] M. E. Cates and C. Nardini, Active phase separation: New phenomenology from non-equilibrium physics, Rep. Prog. Phys. 88, 056601 (2025).

[32] D. Hnisz, B. J. Abraham, T. I. Lee, A. Lau, V. Saint-André, A. A. Sigova, H. A. Hoke, and R. A. Young, Super-Enhancers in the Control of Cell Identity and Disease, Cell 155, 934 (2013).

[33] B. N. Devaiah, A. Gegonne, and D. S. Singer, Bromodomain 4: A cellular Swiss army knife, J. Leukoc. Biol. 100, 679 (2016).

[34] X. Lu, X. Zhu, Y. Li, M. Liu, B. Yu, Y. Wang, M. Rao, H. Yang, K. Zhou, Y. Wang, Y. Chen, M. Chen, S. Zhuang, L.-F. Chen, R. Liu, and R. Chen, Multiple P-TEFbs cooperatively regulate the release of promoter-proximally paused RNA polymerase II, Nucleic Acids Res. 44, 6853 (2016).

[35] B. R. Sabari, A. Dall’Agnese, A. Boija, I. A. Klein, E. L. Coffey, K. Shrinivas, B. J. Abraham, N. M. Hannett, A. V. Zamudio, J. C. Manteiga, C. H. Li, Y. E. Guo, D. S. Day, J. Schuijers, E. Vasile, S. Malik, D. Hnisz, T. I. Lee, I. I. Cisse, R. G. Roeder, P. A. Sharp, A. K. Chakraborty, and R. A. Young, Coactivator condensation at super-enhancers links phase separation and gene control, Science 361, eaar3958 (2018).

[36] X. Han, D. Yu, R. Gu, Y. Jia, Q. Wang, A. Jaganathan, X. Yang, M. Yu, N. Babault, C. Zhao, H. Yi, Q. Zhang, M.-M. Zhou, and L. Zeng, Roles of the BRD4 short isoform in phase separation and active gene transcription, Nat. Struct. Mol. Biol. 27, 333 (2020).

[37] A. W. Fritsch, J. M. Iglesias-Artola, and A. A. Hyman, inPhase — A simple, accurate and fast approach to determine phase diagrams of protein condensates (2024).

[38] S. Alberti, A. Gladfelter, and T. Mittag, Considerations and Challenges in Studying Liquid-Liquid Phase Separation and Biomolecular Condensates, Cell 176, 419 (2019).

[39] W.-K. Cho, J.-H. Spille, M. Hecht, C. Lee, C. Li, V. Grube, and I. I. Cisse, Mediator and RNA polymerase II clusters associate in transcription-dependent condensates, Science 361, 412 (2018).

[40] A. Reicher, J. Reiniš, M. Ciobanu, P. Růžčka, M. Malik, M. Siklos, V. Kartysh, T. Tomek, A. Koren, A. F. Rendeiro, and S. Kubicek, Pooled multicolour tagging for visualizing subcellular protein dynamics, Nat. Cell Biol. 26, 745 (2024).

[41] P. C. Hohenberg and B. I. Halperin, Theory of dynamic critical phenomena, Rev. Mod. Phys. 49, 435 (1977).

[42] K. Hertäg J. F. Robinson, and T. Speck, Statistics and morphologies of stable droplets in scalar active fluids, Phys. Rev. E 113, 015410 (2026).

[43] J. Bauermann, G. Bartolucci, C. A. Weber, and F. Jülicher, Theory of Reversed Ripening in Active Phase Separating Systems, Phys. Rev. Lett. 135, 148201 (2025).

[44] T. Speck, Coexistence of active Brownian disks: Van der Waals theory and analytical results, Phys. Rev. E 103, 012607 (2021).

[45] G. Fausti, M. E. Cates, and C. Nardini, Statistical properties of microphase and bubbly phase-separated active fluids, Phys. Rev. E 110, L042103 (2024).

[46] M. Cates, Dynamics of living polymers and flexible surfactant micelles : Scaling laws for dilution, J. Phys. France 49, 1593 (1988).

[47] C. A. Schneider, W. S. Rasband, and K. W. Eliceiri, NIH Image to ImageJ: 25 years of image analysis, Nat. Methods 9, 671 (2012).

[48] J. Schindelin, I. Arganda-Carreras, E. Frise, V. Kaynig, M. Longair, T. Pietzsch, S. Preibisch, C. Rueden, S. Saalfeld, B. Schmid, J.-Y. Tinevez, D. J. White, V. Hartenstein, K. Eliceiri, P. Tomancak, and A. Cardona, Fiji: An open-source platform for biological-image analysis, Nat. Methods 9, 676 (2012).

[49] G. Koulouras, A. Panagopoulos, M. A. Rapsomaniki, N. N. Giakoumakis, S. Taraviras, and Z. Lygerou, EasyFRAP-web: A web-based tool for the analysis of fluorescence recovery after photobleaching data, Nucleic Acids Res. 46, W467 (2018).

[50] M. Kotecki, Isolation and Characterization of a Near-Haploid Human Cell Line, Exp. Cell Res. 252, 273 (1999).

[51] J. E. Carette, C. P. Guimaraes, M. Varadarajan, A. S. Park, I. Wuethrich, A. Godarova, M. Kotecki, B. H. Cochran, E. Spooner, H. L. Ploegh, and T. R. Brummelkamp, Haploid Genetic Screens in Human Cells Identify Host Factors Used by Pathogens, Science 326, 1231 (2009).

[52] J. F. Robinson, T. Machon, and T. Speck, Universal limiting behavior of reaction-diffusion systems with conservation laws, Phys. Rev. E 111, 065417 (2025).

