## Supplemental Figures for "Active field theory approach to explain size control of transcriptional condensates"

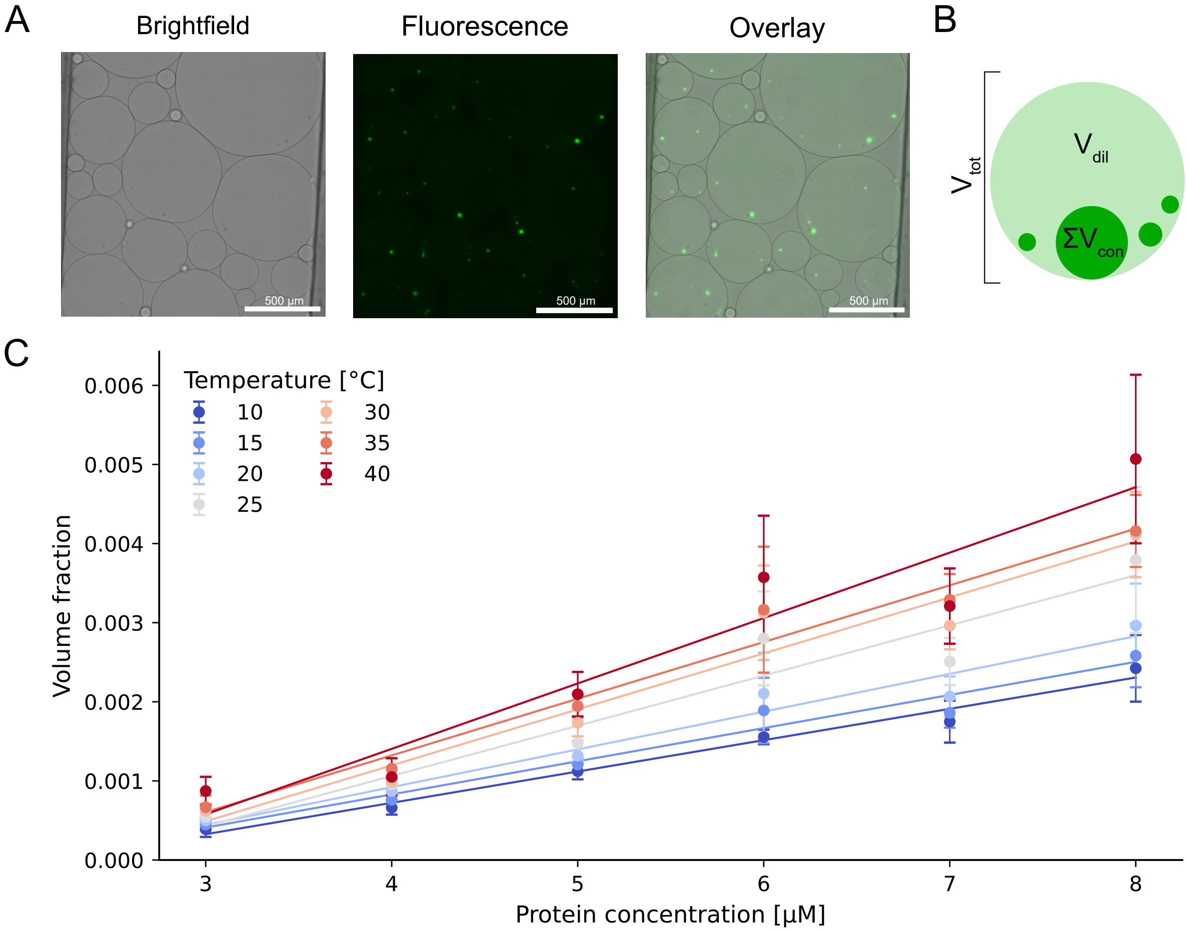


**SUPP. FIGURE 1** | Determination of dense-phase volume fractions using the inPhase assay. (A) Representative microscopy images of BRD4-L emulsions acquired on a temperature-controlled stage, shown as brightfield, fluorescence and overlay (brightfield + fluorescence channels). Fluorescence highlights condensates within individual emulsion droplets (scale bars, 500 µm). (B) Schematic illustrating quantification of the dense-phase volume fraction. For each emulsion droplet, the total droplet volume ($V_{\text{tot}}$) was estimated from droplet geometry and the cumulative condensate volume ($V_{\text{con}}$) was obtained from segmented fluorescence signal. The volume fraction was calculated as $\phi = V_{\text{con}}/V_{\text{tot}}$. (C) Dense-phase volume fraction as a function of total protein concentration for temperatures between 10 °C and 40 °C. Points represent averages of individual droplets, scale bars standard deviation of the mean; solid lines denote linear fits at each temperature.

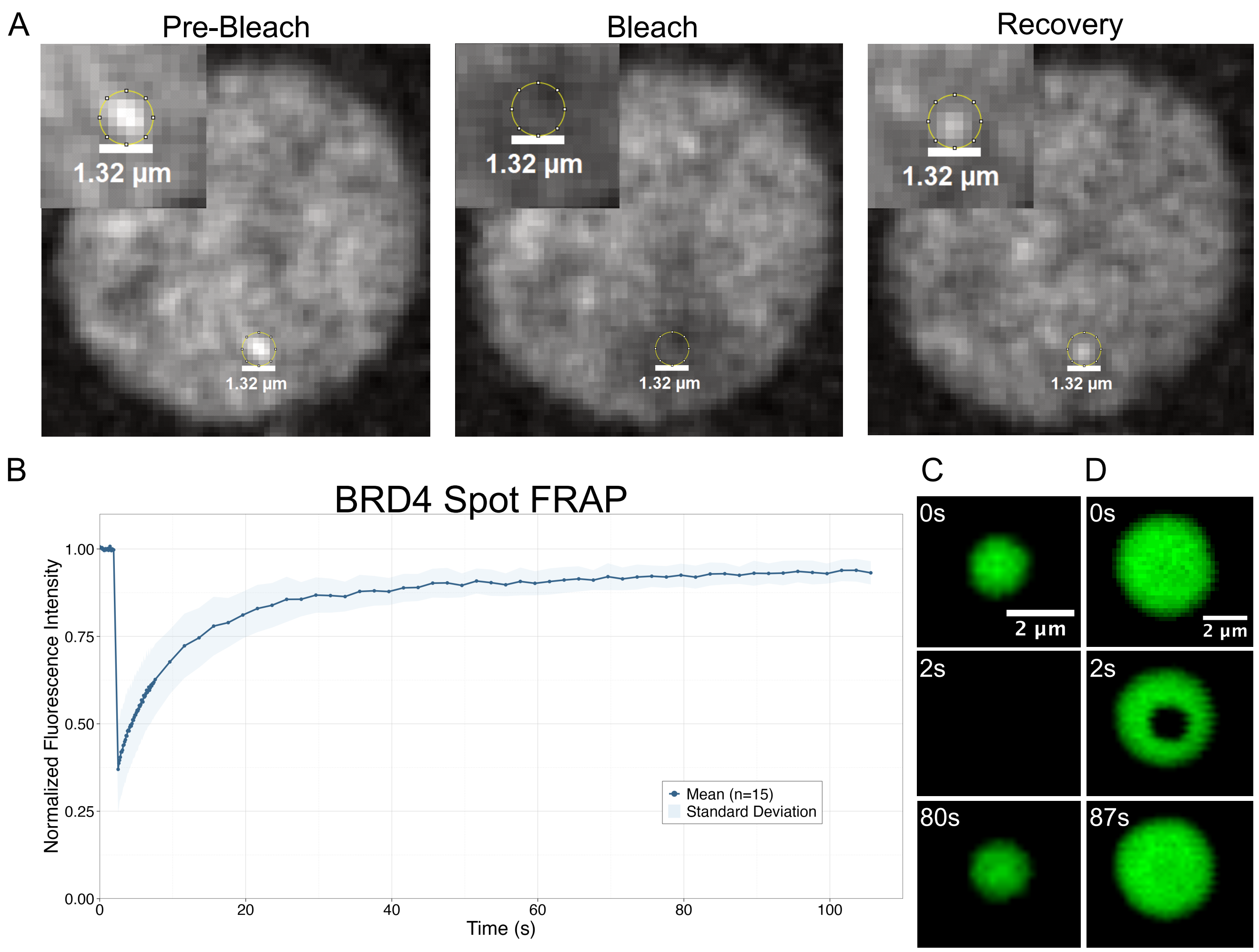


**SUPP. FIGURE 2** | BRD4::eGFP undergoes rapid fluorescence recovery after photobleaching. (A) Representative fluorescence recovery after photobleaching (FRAP) experiment performed on BRD4::eGFP in live HAP1 cells. Confocal images show the bleached region (dashed circle, ∅1.32 µm [6 pixel]) at the pre-bleach (0 s), bleach (+3.2 s), and recovery (+43 s) time points. The recovery curve shows mean relative fluorescence intensity of BRD4::eGFP plotted over time (seconds), fitted with a single-exponential model (fitted maximum relative intensity = 0.86, $R^2$ = 0.86, $t_{1/2}$ = 3.14 s), indicating rapid and near-complete fluorescence recovery consistent with high molecular mobility of BRD4 at transcriptional condensates. (B) Quantification of normalized fluorescence recovery after photobleaching in droplets ($n$ = 15) in vitro (left). Data are represented as mean ±SD in shaded region. (C) Representative images (right) show recovery of fully bleached droplets (column 1) and spot bleached droplets (ROI diameter approx. 1.5 µm; column 2).

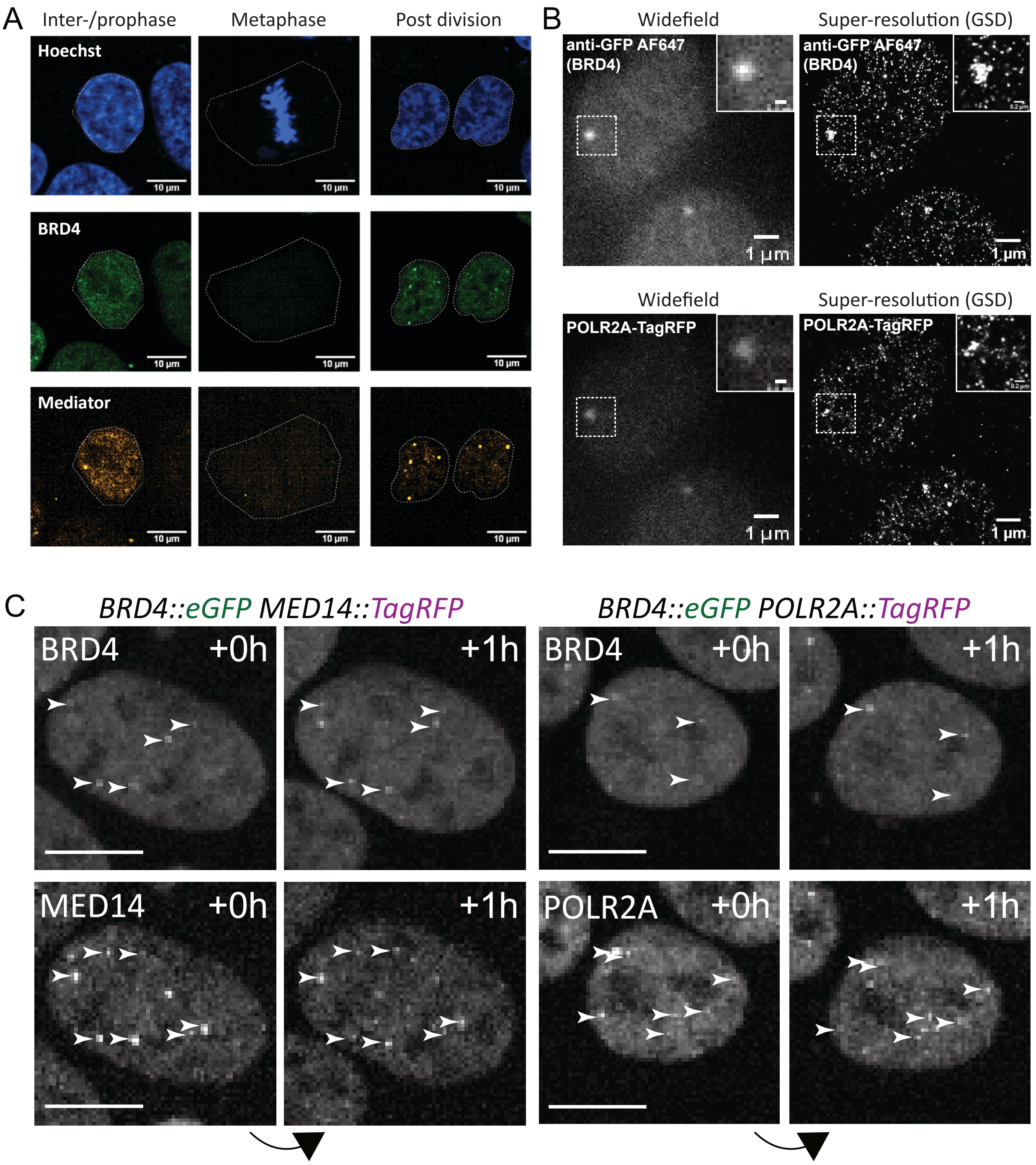


**SUPP. FIGURE 3** | BRD4 and Mediator condensates are dissolved during mitosis and resolved by super-resolution microscopy. (A) Representative live-cell fluorescence microscopy images of HAP1 BRD4::eGFP MED14::TagRFP cells across cell cycle stages. Hoechst (DNA), BRD4, and Mediator (MED14) channels are shown at inter-/prophase, metaphase, and post-division time points (scale bar, 10 µm), illustrating the mitotic dissolution and re-establishment of transcriptional condensates following cell division. (B) Comparison of widefield and super-resolution ground-state depletion (GSD) microscopy images of BRD4 and POLR2A in HAP1 cells. BRD4 was visualized using an anti-GFP::AF647 antibody; POLR2A was imaged via TagRFP. Single-molecule localization processing was applied to generate super-resolution images (scale bar, 1 µm), revealing the sub-diffraction structure of individual BRD4 and POLR2A foci. (C) Time-lapse fluorescence images of BRD4::eGFP MED14::TagRFP (top) and BRD4::eGFP POLR2A::TagRFP (bottom) HAP1 cells at 0 h and +1 h. Arrowheads indicate BRD4/MED14 and BRD4/POLR2A spots that are visible in both timepoints. Images demonstrate the stability and persistence of transcriptional condensate over a one-hour imaging interval.
